# Integrated 3D Light-Sheet and 2D Multiplex Imaging for Deep Histological Profiling of a Somatic Mouse Glioblastoma Model

**DOI:** 10.1101/2025.10.31.685751

**Authors:** Marie Catherine Tiveron, Nathalie Coré, Kevin Bigott, Yliana Hurriaux Fontana, Maria Caccavalle, Lena Vilvandre, Fabio Al Yassouri, Victoria Schöppel, Silvia Rüberg, Melanie Jungblut, Dominique Figarella-Branger, Aurelie Tchoghandjian, Andreas Bosio, Harold Cremer

## Abstract

Glioblastoma is a devastating brain cancer. Despite intense research, patient survival has not significantly increased over the past decades and efficient treatment is currently not available. Therefore, the fundamental understanding of the disease, based on the development of relevant animal models, combined with the development of efficient tools for their deep analysis, represents a priority. Neural Stem cells in the subventricular zone of the forebrain have been identified as cells of origin for glioblastoma, leading to the development of new somatic lineage models based on in vivo brain electroporation. While such models have been characterized in depths by sequencing approaches, systematic histological analyses are currently scarce. Here we present the multimodal histological characterization of a transgenesis independent somatic glioblastoma model in mice. Using 3D light sheet imaging we demonstrate that the model is highly reproducible, allowing quantitative evaluation of tumor growth over large cohorts. Using multiplex imaging by MICS technology we systematically characterize the cellular landscape and molecular composition of the induced tumors, as well as their micro- and macro-environments, and provide a resource of mouse compatible antibodies for cancer research. Finally, we use the model to show that tissue clearing and 3D light sheet microscopy of whole brains can be combined with subsequent multiplex imaging, allowing deep spatial characterization of the tumor proteome in pre-identified brain regions.

## Introduction

Glioblastoma (GBM) represents the most frequent brain cancer in humans. Effective treatment of this type of tumor is still unavailable and the current post-diagnostic survival is below 20 months. (Stupp et al.; Vollmann-Zwerenz et al., 2020). Despite intense research, patient survival has only marginally improved over the past decades, indicating that the currently used approaches inefficiently target the critical cells or address the relevant mechanisms that underlie the malign properties of transformed cells. A deep and fundamental understanding of glioblastoma development, from earliest stages of transformation in initially normal cells to fully developed tumors, represents a prerequisite. Therefore, relevant and well characterized preclinical models are needed.

The demonstration that Neural Stem Cells (NSCs) in the subventricular zone (SVZ) of the human and rodent forebrain ventricles represent cells of origin(Lee et al., 2018a) provided a new experimental entry point into the study of GBM. Indeed, in vivo electroporation allows the efficient introduction of genetic material into NSCs (Boutin et al., 2008a; Bugeon et al., 2021; Coré et al., 2020) and several studies demonstrated that their manipulation by CRISPR-Cas9 mutagenesis and combined with transgenic approaches led to brain tumors with characteristics of high-grade glioma(Clements et al., 2024; Garcia-Diaz et al., 2023; Lee et al., 2018a; Myers et al., 2024; Yamamoto et al., 2021).

Such somatic models show properties that make them important tools for cancer research. For example, they start with a defined population of NSCs as cells of origin, permitting the study of growth tumors development during early stages. Cancer cell lineages can be forced to express fluorescent proteins, allowing their distinction from the non-cancerous microenvironment and the surrounding healthy parenchyma. Finally, tumors develop in fully immunocompetent animals and therefore cancer/immune system interactions can be studied in a functional context closely mimicking human GBM (Chen et al., 2023). Such approaches have been used to investigate, for example, the role of transcription factors in tumor initiation (Myers et al., 2024), the properties of the tumor margin(Garcia-Diaz et al., 2023) brain infiltration (Li et al., 2025)or the tumoral immune-microenvironment (Chen et al., 2023; Chipman et al., 2023; Kim et al., 2022).

Thus, the potential of somatic NSC mouse models in GBM research is considerable, and their deep characterization at the molecular and cellular levels represents an important pre-requisite for their efficient use. Indeed, cell isolation combined with bulk- or single cell sequencing have been successfully applied to provide insight into the gene expression changes in different mouse GBM contexts (Garcia-Diaz et al., 2023; Myers et al., 2024)(Li et al., 2025). However, systematic histological descriptions of such induced tumors, from early transformation of NSCs to full blown GBM, are currently scarce. Moreover, due to the paucity of mouse specific and well characterized antibodies recognizing antigens on the tumor cells and their micro- and macro environments, information about the molecular and cellular organization of these experimental GBM is currently limited.

Here we present the in depths histological analysis of a new electroporation-based lineage model for GBM in mice. Combining Crispr/Cas9 technology, to mutate the tumor suppressor genes TP53 and PTEN, with PiggyBac recombination to stably express the oncogene EGFRviii, we developed a tumor induction system that is independent of transgenic mouse lines and shows all classical histopathological characteristics of GBM. Using tissue clearing and light sheet fluorescent microscopy (LSFM) we followed tumor development in space and time, providing insight into the dynamics of tumor growth and demonstrating that the model is fast, highly reproducible and quantifiable. Using automated cyclic immunofluorescence analysis (MACSima™ Imaging Cyclic Staining (Kinkhabwala et al., 2022)we performed a systematic spatial proteome analysis of the tumors and their environment, validating the model for preclinical studies and providing a resource of functional antibodies that will be crucial for future work in mice. Importantly, we used this model to demonstrate that 3D light sheet microscopy can be combined with MICS, allowing the molecular analysis of predefined tumor regions.

## Results

### Tumor induction by electroporation

We used an electroporation based two-vector system for the induction of brain tumors (Fig. 1A) through the simultaneous inactivation of TP53 and PTEN and the constitutive activation of the EGFR pathway via EGFRviii overexpression, a combination of mutations that has been associated to the classical subtype of GBM (Verhaak et al., 2010). To achieve this, TP53/PTEN gene-inactivation is driven by the CRISPR/Cas9 system (Lee et al., 2018a) while the PiggyBac transposase (PBase) system (Yusa et al., 2009) mediates genomic integration of a second plasmid that assures permanent high-level expression of EGFRvIII and mCherry in all targeted cells and their lineages. Co-electroporation of both plasmids into NSCs along the walls of the forebrain ventricles induced in all animals the appearance of large mCherry positive tumors that extended into neighboring structures of the ipsilateral hemisphere, including the striatum (St), the corpus callosum (cc) and the septum (Sp; Fig. 1B). Also, in most brains at this stage, fluorescent cells were observed in the contralateral brain hemisphere, in a pattern indicating the corpus callosum as a main infiltration route (arrowhead). Analyzing the survival of animals after tumor induction showed onset of morbidity and mortality, in animals that received both plasmids, at about 5 weeks post induction (wpi) and proceeded over the next three weeks (Fig. 1C). Animals that received only a control plasmid or were transfected with the EGFRviii expressing plasmid and PBase alone, showed no lethality over the analyzed time span.

**Figure 1:**
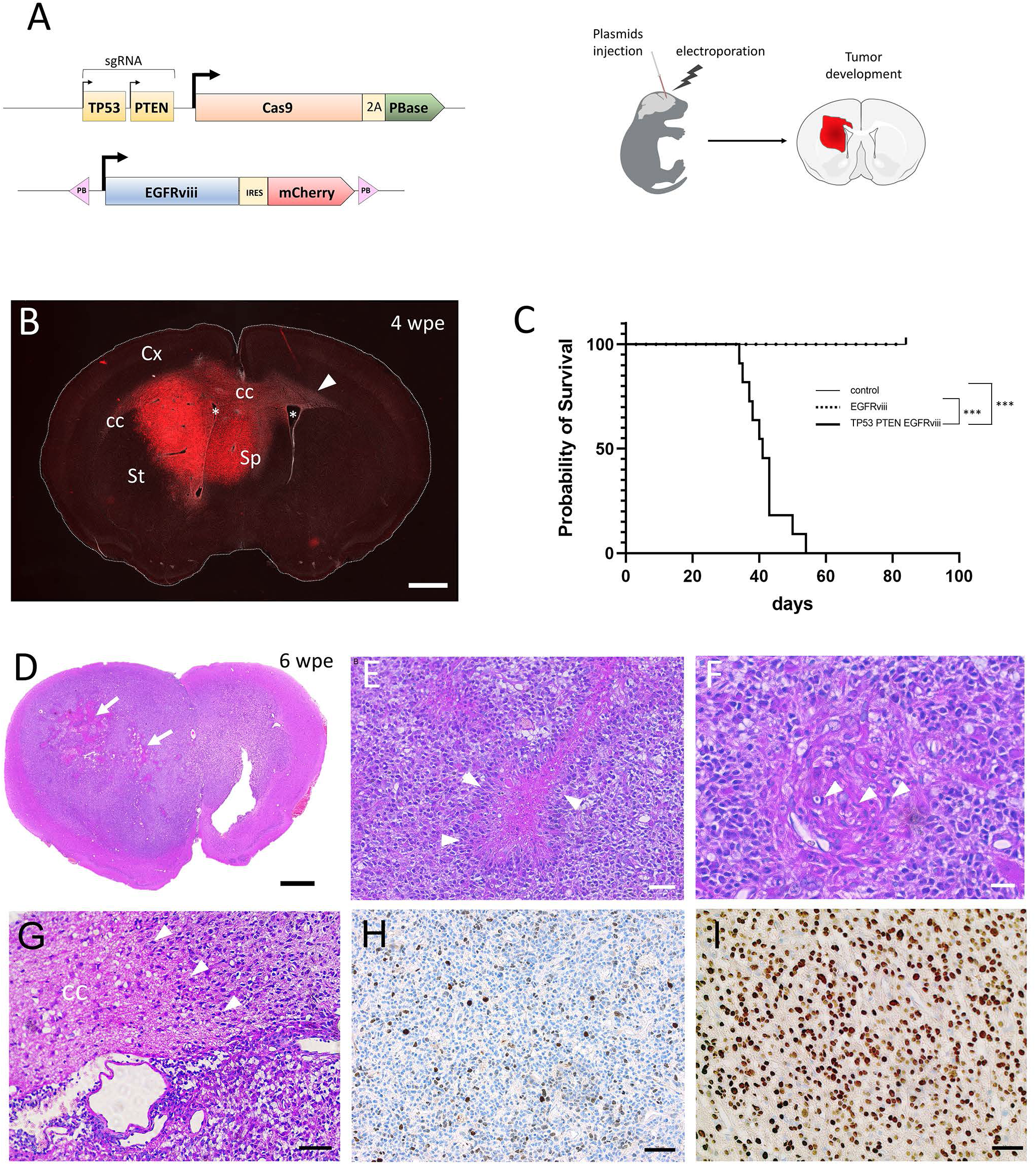
Induction of glioblastoma using plasmid electroporation. (A) Schematic representation of the plasmids used in electroporation. One episomal plasmid carries the CRISPR/Cas9 system mediating the inactivation of TP53/PTEN and supports PiggyBac transposase expression that allows genomic integration of EGFRviii and nuclear mCherry genes encoded by the second plasmid. U6 promoters are used for the expression of each guide RNA while Cas9 and hyPBase are transcribed together under the control of Cbh promoter. EGFRviii and mCherry are expressed from a single transcription unit. After electroporation, only cells targeted with both plasmids will express mCherry and produce tumor formation (small arrows: U6 promoter; large arrows: Cbh promoter) Sg RNA: guide RNA; hyPBase: Hyper PiggyBac transposase; PB: piggyBac elements; P2A: P2A auto-cleaving peptide; IRES: Internal Ribosome Entry Site. (B) Brain coronal section at 4 weeks post electroporation (4wpe). Stars indicate lateral ventricles; arrow head shows tumor cells invading the contra lateral brain hemisphere via the corpus callosum. (C) Kaplan-Meier survival analyses of control (n=10), EGFRviii (n=12) and TP53/PTEN/EGFRviii (n=11) mice. ***: p<0.00001. (D-I) Histological analyses on a brain coronal section at 6 wpe. (D-G) Hematoxylin-Eosin staining with high magnification panels showing necrotic palisade highlighted by arrow heads (E), glomerular vessels with mitotic figures (arrowheads) in endothelial cell (F) and tumor cells invading the corpus callosum (G), arrowheads highlight the migration front). (H-I) Immunohistochemistry for KI67 (H) and Olig2 (I). cc: corpus callosum; Cx: cortex; Sp: septum; St: striatum. Scale bars: 1000 µm (B, D); 50 µm (E, G, H, I); 20 µm (F).

We used a classical histopathological approach to characterize the induced tumors. Representative sections of electroporated animals at 6 wpi, thus with advanced tumors, were analyzed. At this late tumor stage both hemispheres were heavily infiltrated and large tissue lesions were evident (Fig. 1D, arrow). Higher magnification demonstrated the presence of necrotic palisades (Fig. 1E, arrowheads) and glomerular vessels with endothelio-capillar proliferation, representing a hallmark of angiogenesis (Fig.1F; mitoses marked by white arrows), throughout the tumor. In several animals a defined invasion front was identifiable within the corpus callosum (CC) (Fig.1G, arrows), in agreement with previous observations that white matter tracts represents main infiltration route for tumor cells (Claes et al., 2007; Tamura et al., 2019). Finally, immunohistochemistry for the proliferation marker KI67 and the glial marker Olig2 showed high density of positive cells in the tumors (Fig.1HI). Thus, the two-vector lineage model is highly efficient and generates tumors showing the major histopathological hallmarks of classical type GBM (Guo et al., 2021)

### LSFM: Tumor growth in space and time

We next aimed at systematically describing and quantifying GBM growth over time. To appreciate the full spatial complexity of the tumors we combined tissue clearing with light sheet fluorescent microscopy (LSFM), allowing the analysis of tumor size and shape in 3D and in entire brains. We electroporated the GBM induction system either in the lateral (Lateral Wall Electroporation, LWE) dorsal (DWE) or medial (MWE) ventricular wall, as described before (De Chevigny et al., 2012; Tiveron et al., 2017) (Bugeon et al., 2021). For tissue clearing we used the advanced “Clear, Unobstructed Brain Imaging Cocktails” (CUBIC) protocol (Susaki et al., 2014), that allows preservation of fluorescence activity of the introduced marker proteins, here mCherry (Fig. 1A).

Electroporation led to the appearance of small mCherry positive clusters along the ventricular wall at 7 days post induction (dpi, Fig. 2A; Movie 1), in agreement with individually targeted NSCs as cells of origin (Boutin et al., 2008a; De Chevigny et al., 2012). At 2 wpi, individual clusters became indistinguishable. Instead, large mCherry positive cell masses covering the wall and extending into neighboring structures were identifiable, suggesting the fusion of individual clones in a common tumor (Fig. 2B; Movie 2). Over the following two weeks, tumors grew continuously, extending into the surrounding tissues, including the OB and the cortex (Fig.2CD; movies 3,4). DWE and LWE induced tumors presented comparable growths and invasion patterns over time (examples for 2 wpi Fig. S1AB).

**Figure 2:**
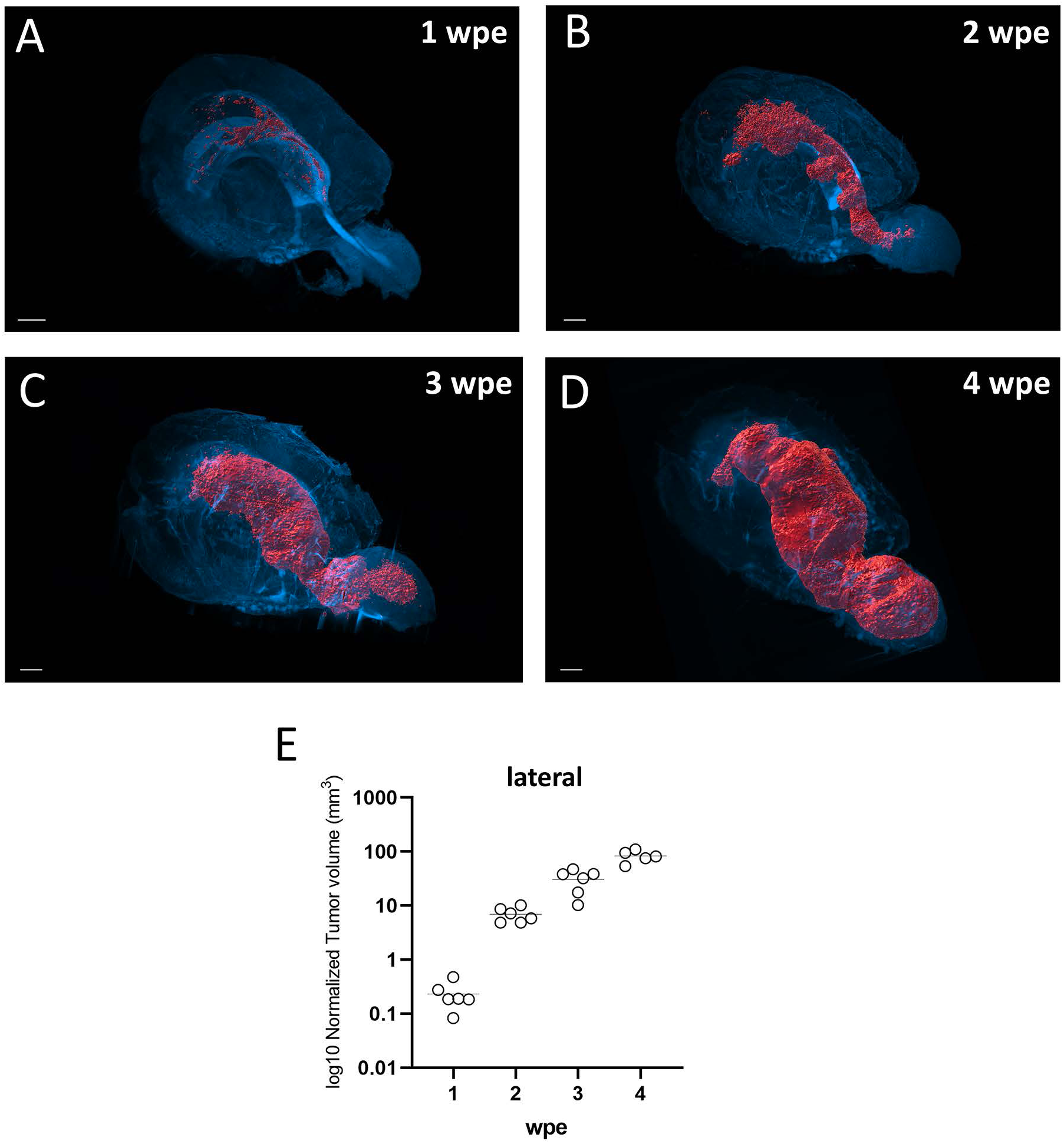
Visualization of tumor progression by 3D imaging. (A-D) LSFM imaging of tumors at 1, 2, 3 and 4 wpe after targeting the lateral wall of the lateral ventricle (LWE). Corresponding 3D animations under Movies 1-4. Segmented signal of mCherry+ tumor cells (red). Nuclei staining with TOPRO3 (blue) allows positioning of the tumor in brain. Note the cell dense SVZ lining the brain ventricles and the rostral migratory stream representing the rostral extension of the SVZ into the olfactory bulb. (E) Growth of tumor volume at 1, 2, 3 and 4 wpe in animals after LWE. Each circle represents one animal. The mean volume per time point per condition is drawn as a bar. Scale bars= 1000 µm.

Quantification of tumor size revealed that growth among animals was highly reproducible over the analyzed period (Fig. 2E, Fig.S1CD). Of note, tumor development, and consequently lethality (Fig. 1C), in this model was very rapid in comparison to other electroporation-based models (Garcia-Diaz et al., 2023; Myers et al., 2024; Yamamoto et al., 2021) likely due to high expression levels of EGFRviii under the control of the Cbh promoter, as well as the genomic integration of multiple copies by the PiggyBac system.

### Multiplex imaging analyses by MICS

To provide deep insight into the histological landscape and molecular composition of the induced tumors, as well as their micro- and macro-environments, we used ultrahigh content imaging based on MICS technology, studying tumors induced in the different ventricular walls at 2 wpi, when compact GBM were observed with full penetrance (Fig. 2B, Fig. S1AB).

Initially, a total of 77 fluorochrome-conjugated antibodies, most of these validated for human tissues but not in mice, were tested on 5 different sections prepared from brains containing tumors induced by either LWE, DWE or MWE. Of these, 38 markers showed high quality staining at cellular level and were analyzed in depths in subsequent steps (Table 1., individual images Fig. S2).

Advanced segmentation using the MACSiQ software (Miltenyi Biotech) identified a total of 20037 cells on a coronal section through an LWE induced tumor at 2 wpi (Fig. 3A). 6206 of these cells were positive for mCherry and 13831 were mCherry negative (Fig. 3B). After-Z score normalization of signal intensities followed by K-d tree analysis and grouping into 5 clusters, 4 clusters comprised mCherry positive cells as well as varying levels of tumor associated proteins, including CD44, GFAP and Nestin (Fig. 3B). In contrast, the cluster comprising cells negative for mCherry contained neuronal markers like Tyrosine Hydroxylase (TH), NeuN and neurofilament, as well as the oligodendrocyte markers Myelin Basic Protein (MBP), MOG and O4, thus representing the surrounding brain parenchyma (Fig. 3BC).

**Figure 3:**
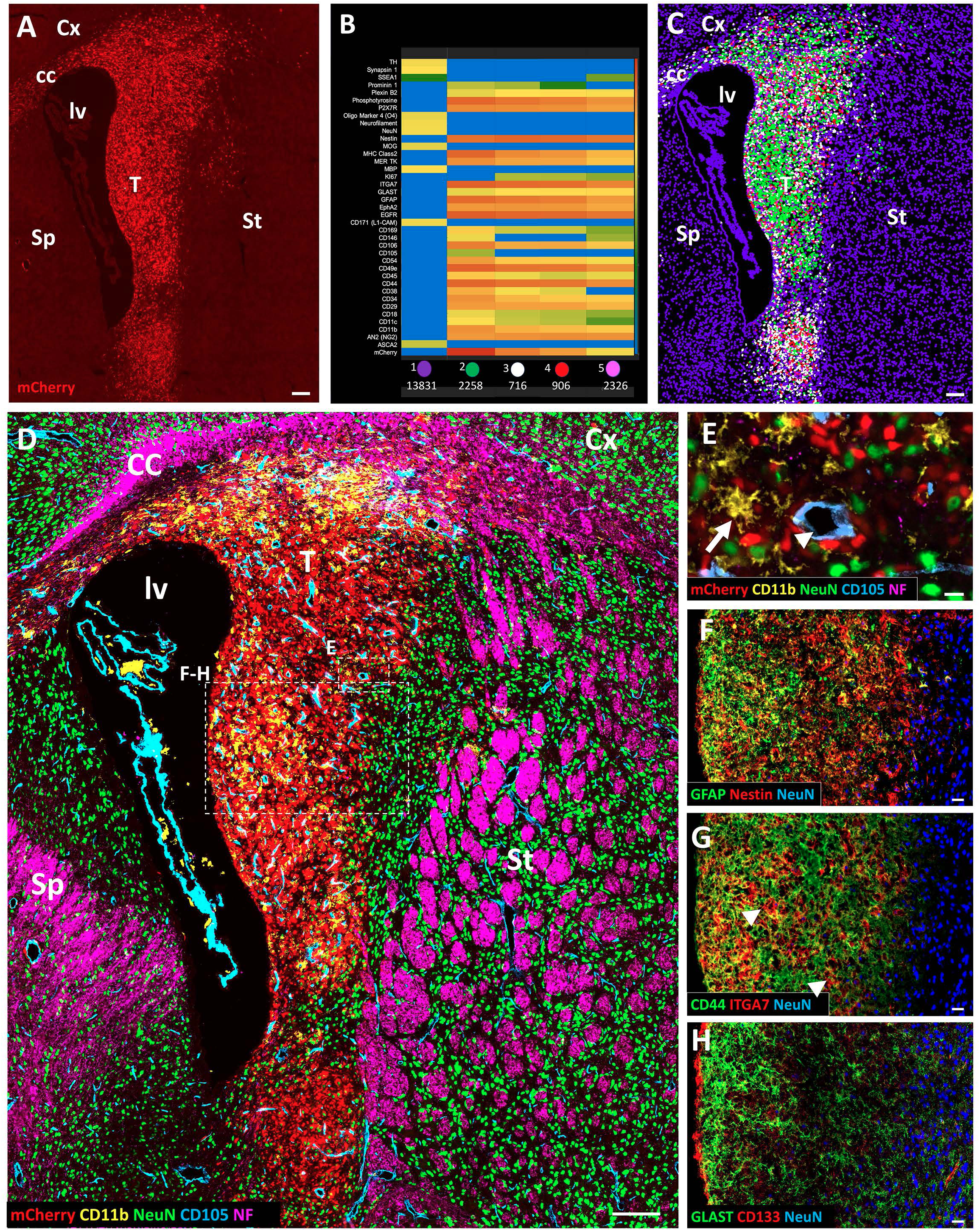
High resolution histological analyses of a tumor generated after 2wpe using MICS. (A) Coronal section used for MICS analyses showing a tumor (T) obtained from LWE at 2wpe. Nuclear mCherry allows identification of tumor cells. (B) Heat map representation of antibody staining after K-d tree analysis and grouping into 5 clusters. One cluster was mCherry negative and comprised neuronal markers like Tyrosine Hydroxylase (TH), NeuN and neurofilament, as well as the oligodendrocyte markers Myelin Basic Protein (MBP), MOG and O4. Four mCherry positive clusters represented tumor cells expressing typical markers like CD44, GFAP and Nestin. (C) Color code representation of cells belonging the 5 clusters presented in B. (D-H) Ultrahigh-content imaging allows a detailed histological overview of the tumor and its microenvironment. (D) Selected markers for tumor cells (mCherry), TAMs (CD11b), vascular endothelial cells (CD105), neurons (NeuN) and axonal tracts (NF; neurofilament). Boxed areas are presented at high magnification in E-H. (E) Example of high-resolution histology obtained with MICS showing CD11b positive TAM (arrow) and a CD105 labeled blood vessel (arrowhead) in the middle of mCherry positive cancer cells. (F-H) Classical tumor markers (GFAP, Nestin, CD44, ITGA7, GLAST, CD133) show labelling in the tumor. Scale bars = 100 µm (A, C, D), 50 µm (F-H) and 5 µm (E). cc: corpus callosum; Cx: cortex; lv: lateral ventricle Sp: septum; St: striatum.

We combined individual markers to provide a detailed histological overview of the tumor, as well as its micro- and macro-environments. We used mCherry (red) for cancer cells, CD11b (yellow) for tumor associated microglia/macrophages (TAMs), CD105 (blue) for vascular endothelial cells, NeuN (green) for neurons and neurofilament (magenta) for axonal projections (Fig. 3D; high magnification of field E in Fig. 3E). All main cell types were easily distinguishable, with a clear separation and complementarity between mCherry positive cancer cell nuclei and NeuN positive neuronal nuclei. Also, strong accumulation of TAMs and large diameter blood vessels was evident within the tumor area (Fig. 3DE). Analysis of coronal sections through tumors induced by DWE and MWE provided images of comparable complexity and quality (Fig. S3A-D).

Next, we focused on the presence of classical tumor markers. In agreement with the astrocytic origin of the induced tumors from postnatal NSCs (Andromidas et al., 2021), GFAP was strongly expressed in tumor regions proximal to the LV, the site of origin of transformed cells. Staining was weaker in distal regions indicating the occurrence of more dedifferentiated and invasive phenotypes. In agreement, Nestin, a marker for undifferentiated brain cells and associated with higher grade gliomas and lower patient survival (Ludwig & Kornblum, 2017; Bradshaw et al., 2016), showed strong and homogeneous staining throughout the entire induced tumor including its invasive margin (Fig. 3F).

Comparably, CD44 was widely and homogeneously expressed over the entire tumor area, in line with its constitutive expression on human tumor cell lines and human GBM (Eibl et al., 1995) (Fig. 3G). In contrast, strongest expression of ITGA7 was confined to defined groups of tumor cells, in agreement with previous findings that identified the integrin as functional marker for glioblastoma stem cells (Haas et al., 2017).

CD133/prominin-1 was strongly expressed on mCherry negative cells lining the ventricular wall, in agreement with its well described expression by a subpopulation of ciliated ependymal cells with NSCs properties (Fig. 3H, arrows; (Coskun et al., 2008)). Within the tumor, merely homogeneous low-level CD133 staining with a punctate aspect was observable (Fig. 3H). Finally, the astrocytic glutamate aspartate transporter GLAST(Corbetta et al., 2019) showed strongest expression in ventricle-proximal regions of the tumor (Fig. 3H), comparable to the expression of GFAP (Fig. 3F).

Next, we investigated the tumor microenvironment (Fig.4A-E). As shown above, CD11b+ TAMs were highly represented within the tumor (Fig.4AB, compare Fig. 3D), generally co-expressed with the leukocyte common antigen CD45 (Fig.4B’). Moreover, within the CD11b/CD45 high population, a large TAM subset expressed CD169 (Fig.4B’’), representing invading blood monocyte derived macrophages (Kim et al., 2022). Finally, a smaller subpopulation of CD11b/CD45 high cells showed labelling for CD11c (Fig.4B’’’), a marker expressed on a broad range of myeloid cells, including activated microglia and dentritic cells (Ricard et al., 2016).

**Figure 4:**
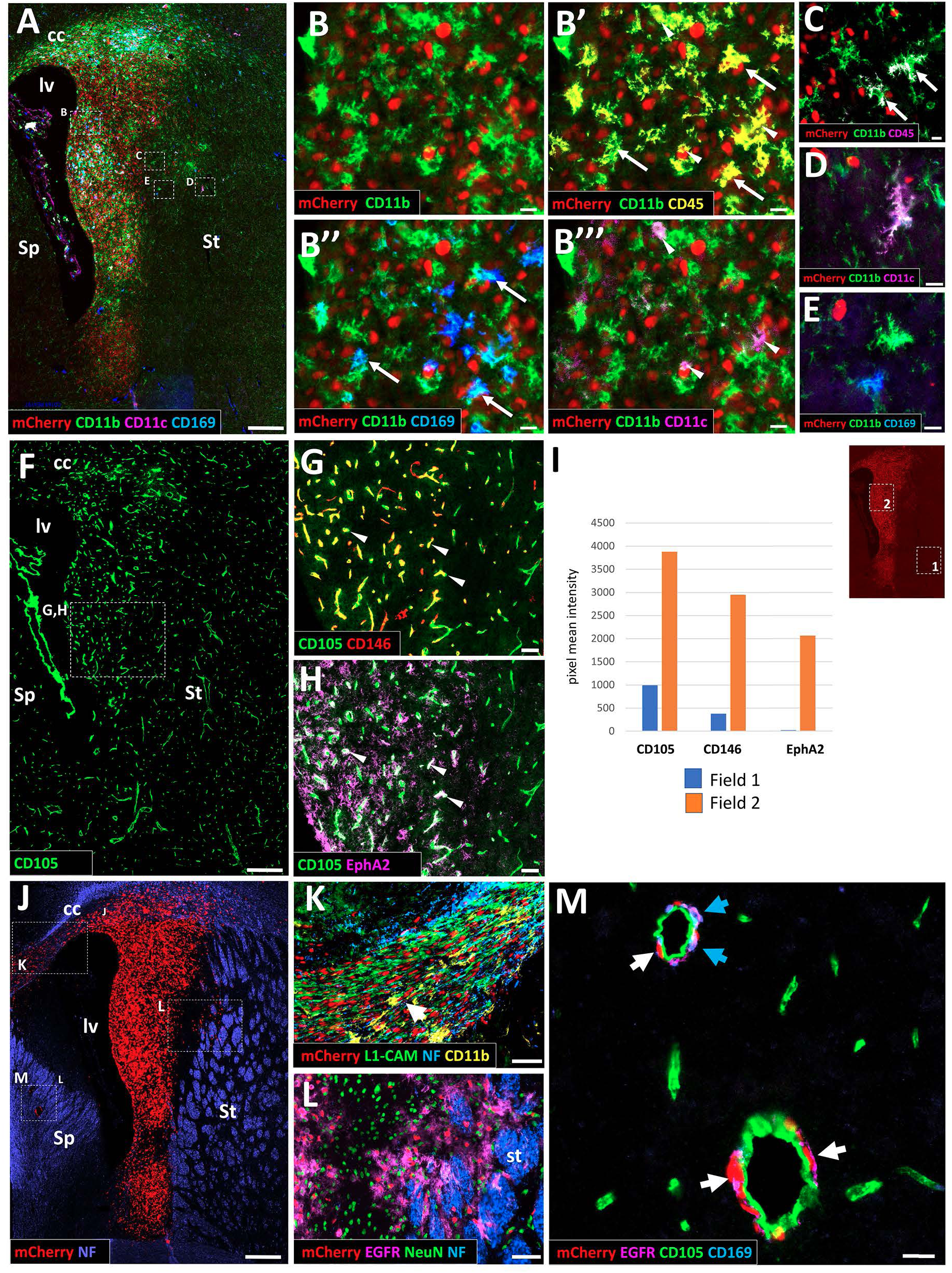
High resolution histological analyses of GBM hallmarks using MICS. (A-E) Tumor immune microenvironment (A) Overview of the section labeled by mCherry (cancer cells), CD11b, CD11c and CD169. Boxed areas delineate regions shown at high magnification in B-E. (B) High magnification of a field of view within the tumor. CD11b positive cells generally co-express CD45, a marker of activated TAMs (B’). Exclusive subpopulations of TAMS are identified with CD169 (arrows, B”) and with CD11c (arrowheads, B’’’). (C-D) High magnification of fields of view at the periphery of the core tumor, where individual tumor cells invade the parenchyma. Activated microglia/TAMs are identified by CD11b and CD45 co-labelling in C (white signal; arrows). TAMs in these peripheral areas are brain resident (D) but also bone marrow-derived (E). (F-I) Tumor associated angiogenesis. (F) Expression of the endothelial marker CD105 reveals higher vasculature density within the tumor. (G and H) High magnifications of marked square in F. (G) Other than the homogeneous expression of CD105, CD146 shows stronger expression in tumor associated blood vessels. (H) EphA2 is confined to the tumor with expression on CD146 positive endothelial cells (arrowheads,) and diffuse presence in the surrounding tumor tissue. (I) Quantification of staining intensity for different vessels markers associated with cancer angiogenesis, outside (field 1) and inside (field 2) the tumor. (J-M) Tumor cell invasion. (J) General view of the tumor (mCherry) surrounded by fiber tracts identified by neurofilament (NF) staining. Boxed areas delineate fields of view presented at high magnification in K-M. (K-L) Migration along neural fibers. In the corpus callosum (K), tumor cells (mCherry) are aligned with L1CAM and NF positive axonal fibers. Some CD11b positive TAMs are also present. At the margin of the core tumor (L), mCherry/EGFR expressing cancer cells are associated with axon bundles of the striatum identified by NF labeling. (M) Migration along the vasculature. In the septum, individual mCherry/EGFR cancer cells are in tight association with CD105 expressing vessel endothelial cells (white arrows) and accompanied by CD169 expressing peripheral TAMs (blue arrows). Scale bars = 250 µm (A, F, J), 100 µm (G, H), 50 µm (K, L), 20 µm (M), 10 µm (B-B’’’), 5 µm (C, D, E). cc: corpus callosum; lv: lateral ventricle, Sp: Septum; St: Striatum.

In addition to TAMs in the core of the induced tumors, macrophages were observed at the tumor margin, in regions were individual mCherry positive cancer cells invaded the surrounding brain parenchyma (Fig. 4 C-E). Here, the presence of TAMS, CD11b-only as well as CD11b/CD45 double positive, indicated early detection of invading tumor cells by immune cells (Fig. 4C). Other distinguishable TAM populations in the infiltration area comprised CD11b/CD11c double positive brain macrophages (Fig. 4D) or CD11b/CD169 bone marrow derived TAMs showing that not only resident immune cells are present at early tumor development stage, but also monocyte-derived cells invading from the periphery. Thus, MICS analyses demonstrated that main populations of TAMs, that have been implicated in human GBM, were identifiable in the induced mouse GBM model.

Then we investigated the expression of markers associated to angiogenesis, another hallmark of GBM. CD105 (Endoglin) was strongly expressed on blood vessels throughout the brain, with significantly higher expression levels in the tumor area versus non tumoral brain tissue, in agreement with the expected high degree of neo-angiogenesis (Fig. 4F-H) (DUFF et al., 2003; Jiang et al., 2012). Expression of CD146 was largely overlapping with CD105 within the tumor (Fig. 4G, arrowheads) but expressed at considerably lower levels on blood vessels of the surrounding normal brain parenchyma (Fig. 4GI), pointing to the well described implication of this VEGFR2 co-receptor in tumor neo-angiogenesis (Zeng et al., 2014). EphA2 has been described as an angiogenic signal in various solid tumors, expressed in both, the cancer cells themselves and the newly generated endothelia (Ogawa et al., 2000). In support of these findings, we observed the presence of EphA2 on CD105/CD146 double positive blood vessels within the tumor (Fig. 4GH, arrowheads) as well as diffuse staining in the surrounding GBM tissue (Fig. 4H). As expected, EphA2 was quasi absent from normal brain tissue outside the tumor area (Fig. 4I).

Then we used the MICS data to study tumor cell invasion, another important hallmark of GBM. White matter tracts, in particular the corpus callosum, represent the most prominent infiltration routes in GBM (Cuddapah et al., 2014; Scherrer, 1938). MICS analyses identified the presence of a stream of mCherry positive cells emanating from the main tumor mass and integrated in the NF and L1-CAM positive fiber tract of the cc (Fig. 4JK). Note also the presence of several CD11b positive immune cells in the cc migratory path (Fig. 4K, arrow). In addition to this massive stream in the CC, individual cells were observed, leaving the lateral tumor margin and invading the neighboring striatum (Fig. 4L). Interestingly, these mCherry/EGFR positive tumor cells were not evenly distributed. Indeed, they were scarce in the region comprising mainly NeuN positive neurons, but accumulated along and inside the NF positive striatal axon bundles, in further support of a white matter guided mode of tumor cell invasion.

Finally, we observed another typical mode of GBM tissue invasion via the brain vasculature. Indeed, we found that two main CD105 positive arteries, localized in the septum, on the opposing side of the LV, thus in considerable distance from the main tumor (Fig.4J), were populated with mCherry/EGFR positive cancer cells (Fig. 4M, white arrows). It is notable in this context that several CD169 positive bone marrow derived macrophages (Fig. 4M, blue arrows) were also present on the cancer cell covered blood vessels, indicating again early interaction of invasive tumor cells with subsets of TAMs.

Altogether, the combined use of classical histological approaches, light sheet 3D microscopy and ultrahigh content imaging by MICS, provides deep insight into brain cancer development in this mouse model, exhibiting dense and morphologically complex tumors that show widespread invasion into healthy parts of the brain. These combined analyses strongly indicate that the model faithfully reproduces the cellular and molecular processes observed in human GBM and provides a resource for mouse-compatible antibodies.

### Combining light sheet microscopy with MICS

Our 3D light sheet microscopy revealed that the induced mouse GBMs were highly complex and spread widely across the rostral-caudal and medio-lateral axes of the analyzed brains (Fig. 2). In parallel, our molecular studies using MICS showed a high degree of molecular and cellular complexity, which could only be examined on individual tissue sections in standard planes, without considering the full three-dimensional extent and morphological complexity of the induced tumors. Our aim was to reconcile these two aspects by performing high-resolution MICS analyses of brains that had previously been analyzed by tissue clearing and LSFM.

First, we induced experimental GBM by LWE, but expressing cytoplasmic GFP instead of nuclear mCherry. After perfusion at 2 wpi brains were permeabilized, stained with an anti-GFP nanobody conjugate and cleared using the MACS® Deep Clearing Kit (Fig. 5A). 3D images were acquired and surfaces were generated with Imaris analysis software (for detail see STAR-PROTOCOLS-D-25-00517). This approach allowed the identification and imaging of tumors at a quality level comparable to the CUBIC approach (Fig. 5B, 3D-animation in Movie 5; brain surface in green, GBM in red;). The main tumor (T) covered the wall of the lateral ventricle, underlying the CC and extending connected groups of cells into the underlying striatum (Fig. 5B, arrows). In addition, a large group of GFP expressing cells was identified in the ventral aspect of the lateral ventricle, separated from the main tumor (Fig. 5B, arrowhead). Finally, a smaller isolated cluster of GFP-high cells was localized in the striatum (Fig. 5B, asterisk; but see Movie 5).

**Figure 5:**
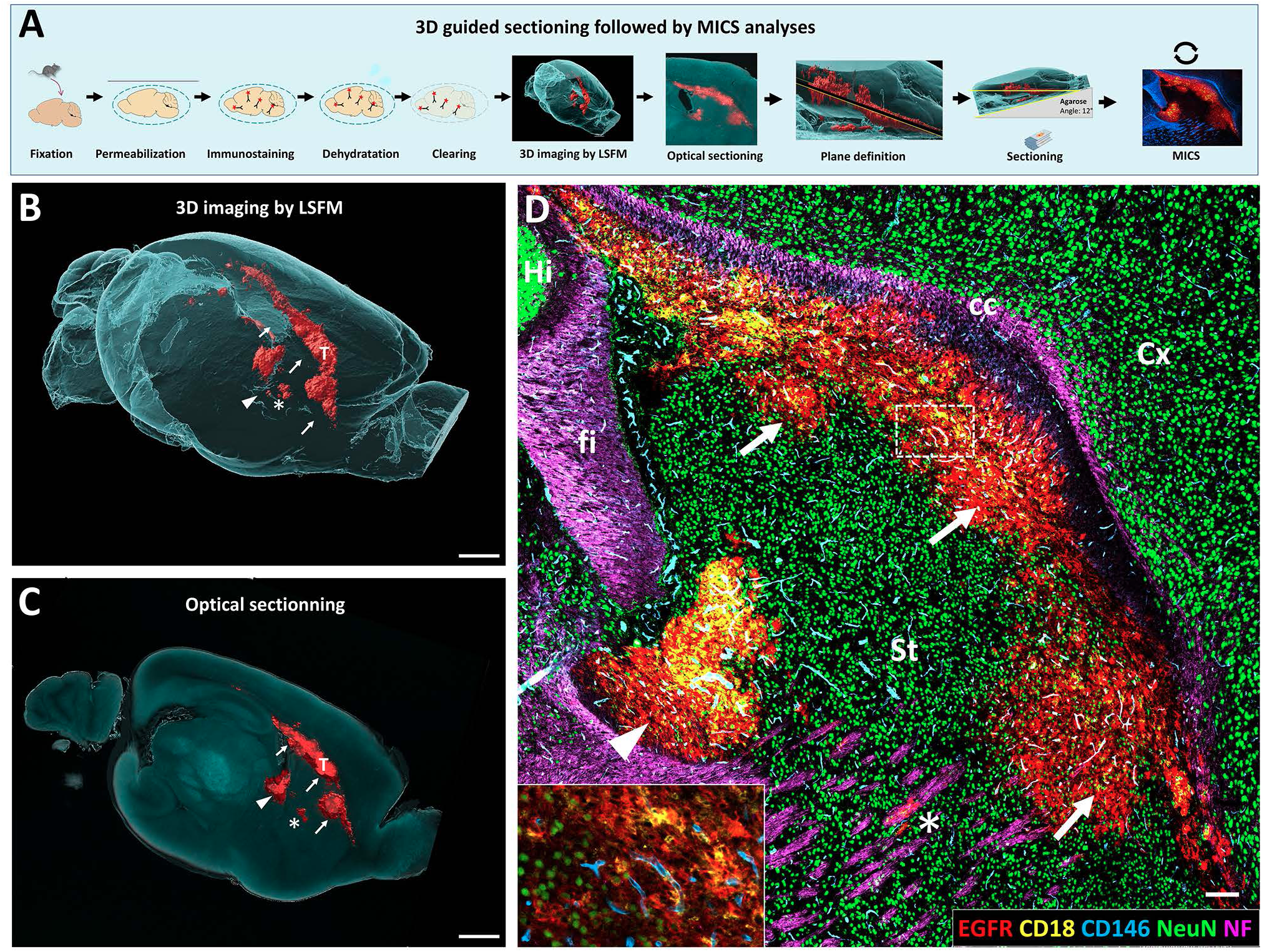
Sequential light sheet microscopy guided sectioning and MICS. (A) Scheme of the protocol used for guided sectioning after imaging by LSFM followed by MICS analyses. (B) 3D image of a brain 2 weeks post electroporation showing the main tumor (T) extending in the neighboring parenchyma (arrows). A small group of cells detached in the striatum (asterisk)and a large non-connected tumor cell mass was observed along the ventral aspect of the ventricle. For animation see Movie 5. (C) Illustration of target plane determination using optical sectioning of the 3D view. (D) High resolution histological analyses using ultrahigh content imaging on a section generated by the 3D-2D protocol. Selected markers showing labeling of the tumor cells (EGFR), leucocytes (CD18), blood vessels (CD146), neurons (NeuN) and axonal fibers tracts (neurofilament; NF). Scale bars: 1000 µm (B, C), 250 µm and 80 µm in inset (D). cc: corpus callosum, Cx: Cortex, fi: fimbria, Hi: Hippocampus, St: Striatum.

Based on these 3D data we used optical sectioning to define an optimal plane that covers the maximum extend of the tumor, including the isolated cell clusters, in a single histological section (Fig. 5C). For the presented brain we defined a 12° deviation from the mid-sagittal plane as optimal and established a protocol allowing precise cryostat sections at this angle (Fig. 5A; STAR-PROTOCOLS-D-25-00517). After re-hydration, cryoprotection, angle correction and snap freezing, a selected 10 μm thick section at the desired level was subjected to MICS. Fourteen of the 38 markers used in the 2D-only MICS approach (Table 1) provided high quality staining (Table 2, individual images in Fig. S4). In addition, the low affinity neurotrophin receptor CD271/p75 (Boiko et al., 2010), that did not provide high quality staining in the MICS-only analyses, showed excellent and specific staining in the 3D2D approach and was further analyzed.

Fig. 5D shows a representative image with superposed staining for tumor cells (EGFR), immune cells (CD18), blood vessels (CD146), neurons (NeuN) and axonal projections (NF). In agreement with the 3D projection (Fig. 5B,C), a main EGFR positive tumor mass was positioned underlying the CC (labelled by NF, magenta) with little to no invasion into the overlying Cortex (NeuN) but extending deep into the striatum (arrows). In addition to the main tumor mass, a large cluster of GFP/EGFR positive cells was detected along the ventral aspect of the lateral ventricle (Fig. 5B,C; arrowheads). Finally, as expected from the LSFM imaging data, a smaller group EGFR positive cells was found separated from the main tumor and integrated in the axon bundles of the striatum, suggesting tissue invasion from the main tumor by a white matter route (Fig. 5D, asterisk). Overall, correspondence between the tissue section and the tumor as detected by LSFM was excellent.

Next, we studied the expression of classical markers for cancer cells in 3D pre-defined regions. Nestin was widely and homogeneously expressed in all parts of the induced tumor (Fig 6A-D). Overlap of Nestin with CD44 expression was strong in regions of the main tumor that were localized in proximity to the CC (Fig. 6C). Extensions of the main tumor into the underlying striatum were also strongly Nestin positive but showed overall lower CD44 expression levels (Fig. 6BD).

**Figure 6:**
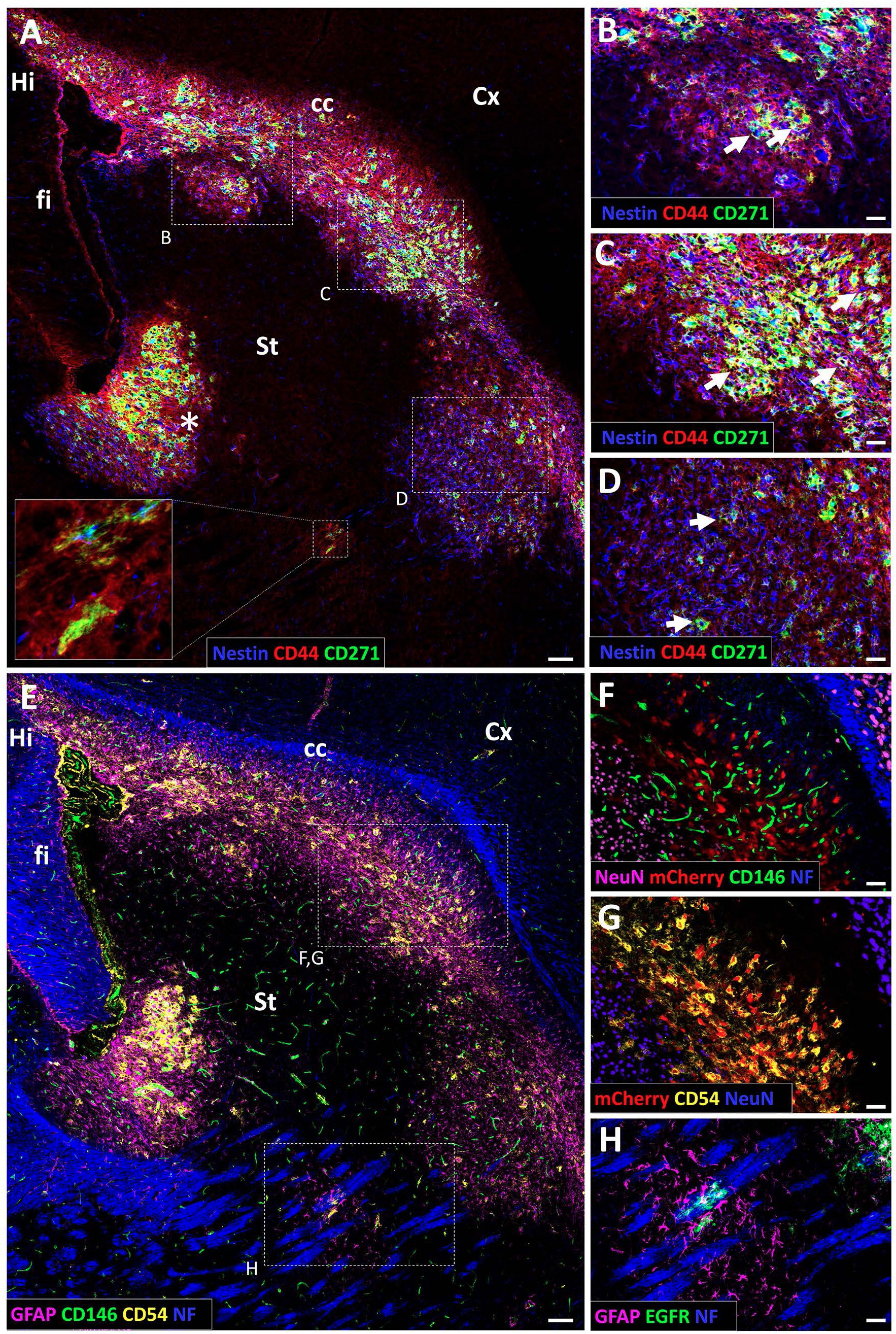
Ultrahigh-content imaging performed after 3D-2D protocol. (A-D) High resolution image showing differential spatial labeling for the tumor markers Nestin, CD44 and CD271. CD271, a marker for cancer stem cell and invading tumor cells, is confined to subpopulations in the main tumor (high magnification in B-D, arrows) and is expressed by invading cells in the striatum (insert in fig. 6A). (E-H) High resolution image showing staining for tumor cells (GFAP and CD54/ICAM1), vasculature (CD146), fibers tracts (NF) and neurons (NeuN). High magnification in F-H. Scale bars= 100 µm (A, E), 50 µm (B-D, F-H).

As mentioned, another tumor marker that provided high quality staining was CD271/p75, a functional marker for tumor-initiating cancer stem cells as well as highly infiltrative cells that emigrate from the main tumor and evade surgical resection (Alshehri et al., 2017). Indeed, high expression levels in humans associate with high-grade glioma and poorer overall survival (Alshehri et al., 2017; Bradshaw et al., 2016). MICS analyses showed expression of CD271 in clusters of Nestin/CD44 positive cells in all regions of the induced tumor (Fig.6A-D, arrows). However, the number of CD271 expressing cells was heterogeneous in different tumor regions, ranging from merely individualized cells in areas extending into the striatum (Fig.6ABD), to a predominance of CD271 positive tumor cells in proximity to the CC (Fig. 6C) and in the group of tumor cells positioned at the ventral aspect of the lateral ventricle (Fig.6A asterisk). Interestingly, CD271 was also expressed in the isolated cluster of cells integrated in the striatal axon bundles (see insert in Fig.6A), further arguing that this group is a result of tissue invasion.

Other examples for markers that showed high quality staining in the 3D2D approach were CD146, CD54/ICAM1 and GFAP (Fig. 6E-H). Indeed, comparable to the 2D-only MICS analyses (Fig.4G), CD146 was a reliable marker of blood vessels in the tumor and at lower levels in the surrounding normal brain (Fig. 6EF). CD54/ICAM-1, a potent cancer cell secreted chemoattractant for immune cells (Zhang et al., 2025) was localized surrounding the mCherry positive nuclei of tumor cells but absent from the normal brain tissue. As expected, GFAP was strongly expressed in the different tumor regions, largely overlapping with the presence of Nestin and CD44 (Fig.6E, compare Fig.6A). Interestingly, strong GFAP positive cells with astroglial morphology were also detected surrounding the above-described separated cluster of EGFR positive tumor cells in the striatum (Fig. 6EH), indicating a rapid inflammatory response of the surrounding brain parenchyma (compare Fig. 4CDE). Altogether, these analyses provide proof-of-principle that tissue clearing and 3D imaging is compatible with the subsequent high content spatial imaging approach provided by MICS, permitting molecular analyses of pre-localized brain regions.

## Discussion

Deep characterization of preclinical models, based on a well-equipped toolbox, represents a prerequisite for their efficient use and is required for the translation of experimental result to clinical applications. Our work presented here combines classical histological approaches with latest spatial proteomic and 3D imaging technology to characterize a relevant electroporation based GBM model in mice.

Combining CRISPR/Cas9 induced mutagenesis of TP53 and PTEN with transposase mediated genomic integration of the oncogene EGFRviii we provide a simple and reliable mouse model that fulfils all histopathological criteria for highest grade glioma. It can be applied in basically all genetic contexts in mice, without the laborious and time-consuming generation of complex allelic combinations.

Observation of induced tumors in their entire extend, based on tissue clarification and 3D LSFM, demonstrates that tumor onset and growth in this model is highly reproducible and measurable over large cohorts and prolonged periods, starting as early as one week after transformation. Using multiplex tissue imaging, we investigate the molecular and cellular composition of induced tumors and provide a list of mouse-compatible antibodies that allow the high-resolution analysis of the induced GBM, as well as their micro- and macroenvironments. Finally, we show that tissue clearing and LSFM analyses are compatible with the subsequent multiplexed spatial proteome analysis by MICS technology, providing a new analytic approach that allows serial 3D to 2D analyses.

Over the past decade a wide spectrum of mouse models for the induction of high-grade gliomas have been developed and used to provide insight into specific aspects of GBM and their interactions with the surrounding healthy tissue (Garcia-Diaz et al., 2023; Kim et al., 2022; Myers et al., 2024) (Chipman et al., 2023; Hettiarachchi and Park, 2025). Several of these models are based on the gain of function of EGFR, a potent driver of neural stem cell proliferation, migration and differentiation, that has been identified as one of the key drivers of GBM (Lee et al., 2018b) and represents an important potential target for CAR-T cell based therapeutic approaches (Sterner and Sterner, 2024). The model described here leads, in contrast to other EGFR based models in which tumor induction and progression occur over weeks and month (Kim et al., 2025), to rapid and homogeneous induction of tumors, likely due to the high levels of EGFRviii expression under the control of the actin-based CBh promoter.

While this rapid induction appears at first sight as a potential disadvantage, compared to models based on more physiological EGFR levels, our histopathological analyses clearly show that the resulting tumors present all key features of GBM. Moreover, its rapidity and reproducibility in tumor induction provide access to the events occurring during the very first stages of NSC transformation. Finally, the homogeneous and quantifiable growth of tumors, that we show in our LSFM studies, and the fact that GBM can be induced in various genetic contexts without the need for complex and time-consuming breeding, has potential to provide an suited means to study the molecular regulation of brain cancer development and the efficient preclinical testing of therapeutic approaches.

An important aspect of our work represents the application of high content imaging for the molecular characterization of induced mouse tumors. While multiplex spatial proteomics approaches based on MICS have been widely used on human tissue (Kinkhabwala et al., 2022; Scheuermann et al., 2024), the use of this technology in GBM research in mice is currently limited (Bastiancich et al., 2024). To our knowledge multiplex spatial imaging has not been applied to somatic mouse models.

One reason for this paucity is likely the lack of sufficient well characterized antibodies that provide high quality staining on rodent tissue. Indeed, most available antibodies recognizing cancer cells, or relevant cells of the micro- and macroenvironments, have been raised and selected for work in humans. In agreement, we found that out of an initial panel of 76 human validated antibodies, only about half (38) showed high quality staining on mouse GBM sections. However, despite this limitation, the here identified antibodies comprised markers for almost all relevant tumoral and non-tumoral cell subpopulations. Typical GBM markers like for example GFAP (Simone et al., 2023), nestin (Ludwig and Kornblum, 2017), CD44 (Inoue et al., 2023) or GLAST (Corbetta et al., 2019) were strongly overexpressed in the tumor baring area, in patterns and cell populations that further confirmed the relevance of our GBM model. Another example is the p75 neurotrophin receptor CD271, that has been functionally implicated in two critical glioma cell populations in human, chemoresistant cancer stem cells and infiltrative cells, that efficiently exit from the main tumor and escape surgical resection therapy (Alshehri et al., 2017). In agreement with these findings we identify in mouse GBM strong CD271 expression in a small subset of cells in the tumor core areas, but also in cells escaping from the tumor by white matter tract guided migration.

A major advantage of somatic glioblastoma mouse models, like the one presented here, is that tumor development occurs in a fully immunocompetent environment. In humans, TAMs represent up to 40% of the entire tumor mass, comprising principally brain-resident microglia and bone marrow-derived myeloid cells invading the brain from the periphery (Buonfiglioli and Hambardzumyan, 2020). In agreement we find that as soon as 2 weeks after induction the induced mouse tumors contain large amounts of cells co-expressing CD11b, a marker for basically all TAMs, and the leukocyte common antigen CD45(Peres et al., 2024). In addition to these general markers, CD11c, a marker of primed microglia and infiltrating monocyte-derived dendritic cells (Ricard et al., 2016), was found on subset of TAMs. Finally, we identified a major immune subpopulation expressing CD169, representing bone marrow derived myeloid-derived cells that contribute to anti-tumor immunity (Kim et al., 2022). Thus, while the set of markers for immune cells used here is far from being comprehensive, it provides an entry point into the deep spatial proteomic analyses of tumor-immune system interactions. Indeed, the prevailing hypothesis concerning tumor-immune system interactions suggests that TAMs are initially recruited to eliminate cancer cells but become reprogrammed by the tumor microenvironment to support tumor growth. However, currently most data supporting such a scenario is based on the analysis of late-stage human tumors or fully established tumors in mouse models, which overlook the early, critical phases of immune involvement (Kloosterman et al., 2024; Ochocka et al., 2021; Van Hove et al., 2019). Combining models enabling the controlled progress of gliomagenesis from earliest stages to lethal tumors, as it is possible with the somatic model described here, with deep molecular phenotyping by spatial proteomics, appears as well suited to advance this therapeutically highly relevant field of tumor-immune interactions.

An important aspect of this work is the combined use of tissue clearing and 3D light sheet imaging with targeted 2D spatial multiplex analysis. Indeed, 3D microscopic approaches allow the detection of a limited marker set in the correct spatial context(Snacel-Fazy et al., 2024). In contrast, 2D multiplexing studies provide deep insight into the molecular landscape of a given tissue. However, targeting a region of interest containing as much information as possible in a single section can be challengingThe workflow described here allows to overcome these limitations and therefore maximizes the recovered spatial information.

Based on high resolution 3D images we were defined an ideal target level that allowed the preparation of pre-defined tissue sections that precisely cover this region, a procedure we termed ‘light sheet guided sectioning’ (STAR-PROTOCOLS-D-25-00517). The generated section comprised not only various regions of the induced tumor, that we were able to analyze and compare in the full 3D context, but also allowed the targeted inclusion of separated invading cells, that would almost certainly have been missed on a random tissue section prepared in a standard plane.

MICS analyses of the predefined tissue sections provided high quality staining for 15 markers. While this list is limited, we were able to identify antibodies labeling both tumor cells and the tumor microenvironment. A systematic study using a wide spectrum of potential markers, and also testing other parameters like for example fixation method or antibody concentration, will like provide a far more comprehensive list of antibodies suited for 3D2D sequential approaches.

## Materials and methods

### Mice

All mouse experiments were approved and performed according the guidelines of French Ethical committee according to the European rules and regulations (Authorization number #34083-2021112215573540v5). Swiss CD1 mice (Charles River, France) were group-housed in regular cages under standard conditions on a 12 hours light-dark cycle. The day of birth was defined as post-natal day 0 (P0). Both male and female were used in all experiments. Mice were monitored daily and euthanized if they began to show signs of disease (hunching, limited activity) and reached humane endpoints. This takes place between 34 and 54 days and only occurs in the GBM induction condition.

### Plasmid construction

The two vectors we used to induce high grade glioma were constructed as follow. The vector that allows the expression of the CRISPR Cas9 system together with the PiggyBac transposase, was generated from phU6-sgTP53-sgPTEN-Cbh-Cas9-Cre (kind gift from Dr. Jeong Ho Lee, KAIST, South Korea) where the Cre sequence was replaced by hyperactive PiggyBac transposase (hyPBase)(Yusa et al., 2011) from pCaggs hyPBase (phyPBase, kind gift from Dr. Karine Loulier). The PiggyBac expression vector, that carries EGFRviii and a fluorescent reporter protein, was obtained from pPB-CMV-TO-EGFRvIII-IRES-nlsCherry (gift from Michael Elowitz, Addgene plasmid # 116039) by replacing CMV promoter with Cbh promoter (Gray et al., 2011). This construct was further modified to replace nlsCherry gene by eGFP sequence to obtain PB-Cbh-EGFRviii-IRES-eGFP (PB-EGFRviii-GFP).

### Postnatal electroporation

*In vivo* electroporation was performed as previously described (Boutin et al., 2008b). Briefly, P0-P1 pups were anesthetized by hypothermia and injection of 1.5 µl of a mixed solution of each plasmid at 1µg/µl were injected in the lateral ventricle. Animals were subjected to five 95V electrical pulses (50 ms, separated by 950 ms intervals) using the CUY21 edit device (Nepagene, Chiba, Japan) and 6 mm tweezer electrodes (CUY650P10, Nepagene) coated with conductive gel (Signagel, Parker Laboratories, USA). To target effectively the dorsal or lateral V-SVZ, the positive electrode was positioned respectively, dorsally above the eyes or slightly ahead of the eyes, and the negative electrode was placed in the opposite position. After electroporation, animals were reanimated in a 37°C incubator before returning to the mother.

### Brain Tissue preparation

Mice were deeply anesthetized with a lethal dose of xylazine and ketamine (200 and 20 mg/kg respectively). Intracardiac perfusion was performed with cold PBS; heparin (5u/ml Sigma-Aldrich), to wash out efficiently blood cells from the vasculature, followed by fixation with cold 4% paraformaldehyde in PBS (32% stock solution, Electron Microscopy Sciences). Brains were then dissected out, post fixed overnight then stored in PBS. For MICS analyses, brains were post fixed for only 2 hours, washed quickly in PBS, then cryoprotected in a 30 % sucrose-PBS solution, embedded in OCT (Tissue-Tek) and finally frozen in isopentane cooled in dry ice. Samples were cryosectioned with a CM3050S cryostat (Leica) and 10 microns coronal sections were mounted on Superfrost glass slides and stored at −80°C until processed. For the combination of light sheet microscopy and MICS, brains were post fixed for only 2 hours, washed quickly in PBS and stored in PBS containing sodium 0.09% azide.

### Immunohistochemistry and histology analyses

For histology, serial 5-μm formalin-fixed paraffin-embedded (FFPE) sections of mouse brains were stained with hematoxylin and eosin (H&E) and examined under the microscope for the presence of tumor cells. Immunohistochemistry was performed on adjacent paraffin sections. After steam-heat-induced antigen retrieval, sections of FFPE samples were tested for the presence of OLIG2 (Clone EP112, Zytomics, dilution 1:100) and Ki67 (Clone Mib-1, DAKO, dilution 1:200). A Benchmark Ventana autostainer (Ventana Medical Systems) was used for detection, and slides were simultaneously immunostained in order to avoid intermanipulation variability. Slides were then scanned (Nanozoomer 2.0-HT, Hamamatsu Photonics SARL France, Massy, France) and images processed in NDP.view2 software (Hamamatsu).

### Statistical analysis

Survival data were analyzed using Kaplan-Meier method. Statistical comparison between groups were performed with Mantel-Cox (log-rank) test using GraphPad Prism version 9.0 (GraphPad Software, USA). All P values less than 0.05 were considered statistically significant.

### Brain clearing and 3D imaging for Brain volume assessment

#### Clearing

Brains were cleared using the advanced Clear, Unobstructed Brain Imaging Cocktails (CUBIC) protocol (Susaki et al., 2014). Fixed hemi brains recovered at 1, 2, 3 and 4 wpe were incubated in 7 ml of CUBIC reagent 1 [25% urea, 25% *N*,*N*,*N*′,*N*′-tetrakis-(2-hydroxypropyl)ethylenediamine, and 15% Triton X-100] at 37°C under gentle shaking until complete clearing. CUBIC reagent 1 solution was changed after the first day of incubation and then every 2 days. When brains were transparent, they were washed for 1 hour in PBS and then incubated in TOPRO-3 iodide nuclear dye solution (1:2000; T3605, Life technologies) overnight at 37°C. Brains were further washed in PBS 2-3 times for 10 min and immersed back in CUBIC reagent 1 for one more day at 37°C.

#### Light Sheet Imaging

Cleared brains were imaged in CUBIC1 solution in horizontal orientation with the Ultra Microscope Blaze^TM^ (Miltenyi Biotec, Germany) equipped with a sCMOS 5.5Mpx Camera, LaVision Bio Tec PLAN 1x/0.1 objective lens fitted with a 16 mm working distance dipping cap and LED lasers at 488, 561 and 639 nm. Emission filters used were 525/50, 595/40, 680/30. Stacks of images were used to perform 3D reconstruction.

#### Image analyses

3D projections were performed using Imaris x64 software (Bitplane, Zurich, Switzerland). The volume of tumors (V_Tumor_) was defined by creating a 3D mask using mCherry labelling and the automatic surface tool from Imaris software. Because CUBIC protocol causes tissue swelling, tumor volume was normalized over the volume of the dentate gyrus (DG), a well-defined anatomical structure. The contours of the DG were drawn manually with the Surface tool using TOPRO3 signaling to generate a 3D mask and define the DG volume (V_DG_). Normalized tumor volume was obtained by dividing V_Tumor_ by V_DG_.

### MICS experiment

#### Tissue preparation for MICS

Perfused brains treated as described above were post fixed for 2 hours only, washed quickly in PBS, cryoprotected in a 30 % sucrose-PBS solution, then embedded in OCT (Tissue-Tek) and finally snap-frozen in isopentane cooled in liquid nitrogen. Samples were cryosectioned with a CM3050S cryostat (Leica) and 10 microns coronal sections were mounted on Superfrost glass slides according to MACSwell™ Imaging Frame template and stored at −80°C until processed with MACSima™ imaging cyclic staining (MICS) technology as described in Kinkhabwala et al. 2022 (Kinkhabwala et al., 2022) and Scheuermann et al. 2024 (Scheuermann et al., 2024).

#### Antibodies and reagents for MICS

Primary fluorochrome-labeled antibodies conjugated to FITC, PE APC were used for MICS. Antibody clone used, their dilution and conjugated fluorochrome are indicated in Table 1.

#### MICS

MACSima™ imaging cyclic staining (MICS) was performed for deep cell phenotyping. Samples were mounted on the selected MACSwell™ Imaging Frame and rehydrated with MACSima™ Running Buffer for 2 minutes. DAPI pre-staining (1 µg/mL) was performed in MACSima™ Running Buffer for 10 minutes and washed three times with MACSima™ Running Buffer. Subsequently, MICS technology was applied with iterative staining, imaging and erasure cycles, enabling ultrahigh-content imaging on a single sample as described in Kinkhabwala et al. 2022 (Kinkhabwala et al., 2022) and Scheuermann et al. 2024).

#### 3D-2D STAR Protocol

Find detailed protocol published in STAR Protocols (STAR-PROTOCOLS-D-25-00517).

#### 3D spatial analysis (LSFM)

Two weeks post electroporation with a phU6-sgTP53-sgPTEN-Cbh-Cas9-Cre-hyPBase/PB-EGFRviii-GFP vector combination, brain hemispheres were stained and cleared according to MACS® Deep Clearing Kit protocol (Miltenyi Biotec, 130-136-470). Briefly, after perfusion, samples were permeabilized for 24 hours in Deep Permeabilization Solution at room temperature, followed by immunostaining for 7 days at 37 °C with an anti-GFP AlexaFluor647 nanobody conjugate (Proteintech/ChromoTek, gb2AF647-50, 1:1000) and washed at room temperature, both in Deep Antibody Staining Solution. Dehydration was performed at 28 °C in an ascending ethanol series in distilled water (30%, 50%, 70%, 90%, 100%, 100%) with added 2% Tween20 (v/v). Tissues were rendered transparent in high refractive index Deep Clearing Solution before imaging with an UltraMicroscope Blaze™. 3D light sheet data was acquired with both 1.1x/0.1 NA/≤17 mm WD and 12x/0.53 NA/≤10.9 mm WD objectives for overview and detail view, respectively. Fast imaging of detailed image stack was enabled by the combination of "Fast Tiling Scan" and "Speed Mode" within ImSpector software (Miltenyi Biotec).

#### 3D to 2D transition

Surfaces for both autofluorescence and anti-GFP Alexa Fluor647 signal (GBM) were created within Imaris analysis software. With these surfaces, an angle between mid-sagittal brain hemisphere surface and an optimal target intersection of the GBM could be determined via two "Oblique Slicer" planes. An agarose block (1.5% low melting agarose) with this specific angle was generated. Rehydration was performed at 28°C in a descending ethanol series in distilled water (100%, 90%, 70%, 50%, 30%) with added 2% Tween20 (v/v), followed by two 1xPBS incubations.

Samples were cryoprotected at 2-8 °C in 30% sucrose-PBS solution until fully sedimented. Excess sucrose-PBS solution was removed and tissue were placed on top of the previously generated angled agarose block to harmonize the cutting surface and the GBM target section. Agarose block and tissue were carefully covered with optimal cutting temperature (OCT) compound within a cryo mold and snap-frozen in liquid nitrogen cooled isopentane. Sample were stored at −80 °C until further processing.

#### 2D sample preparation

Sample were mounted within a cryostat as parallel to the blade as possible to intersect the surface fully, which ultimately preserved harmonization of surface and target plane. Sectioning was conducted at −20 to −23 °C and 8-10 µm slices were collected on SuperFrost® slides with a MACSwell™ frame template. Target region was verified with anatomical landmarks on a stereo microscope as well as by detection of preserved 3D-immunofluorescence signature with a fluorescence microscope.

#### 2D spatial analysis (MICS)

MACSima™ imaging cyclic staining (MICS) was performed as described before. 3D-immunofluorescence signature of GBM was acquired in APC-channel during autofluorescence acquisition.

## Acknowledgements

We thank JH Lee (KAIST, Daejeon, South Korea) and Karine Loulier (INM, Montpellier, France) for CRISPR and pBASE plasmids, respectively. Caroline Blanc is acknowledged for help with experimental work. The Cremer lab has been supported by the ANR (Uncoding, ANR-21-CE16-0034; Miniature, ANR-21-CE13-0003; Goligo, ANR-21-CE16-0023) and the Association pour la Recherche contre le Cancer (ARC). Yliana Hurriaux Fontana was supported by a PhD fellowship from the FRM. We also thank the imaging facility at IBDM, member of the National Infrastructure France-BioImaging (https://ror.org/01y7vt929) supported by the French National Research Agency (ANR-24-INSB-0005 FBI BIOGEN) as well as the animal facilities. The Tchoghandjian lab was supported by ANR (MEGA01, ANR-21-CE18-0047-01), the Cancéropole PACA (PETRA structuring action) and the patient association pour la recherche sur les tumeurs cérébrales (ARTC Sud).

---

**Supplementary S1.**
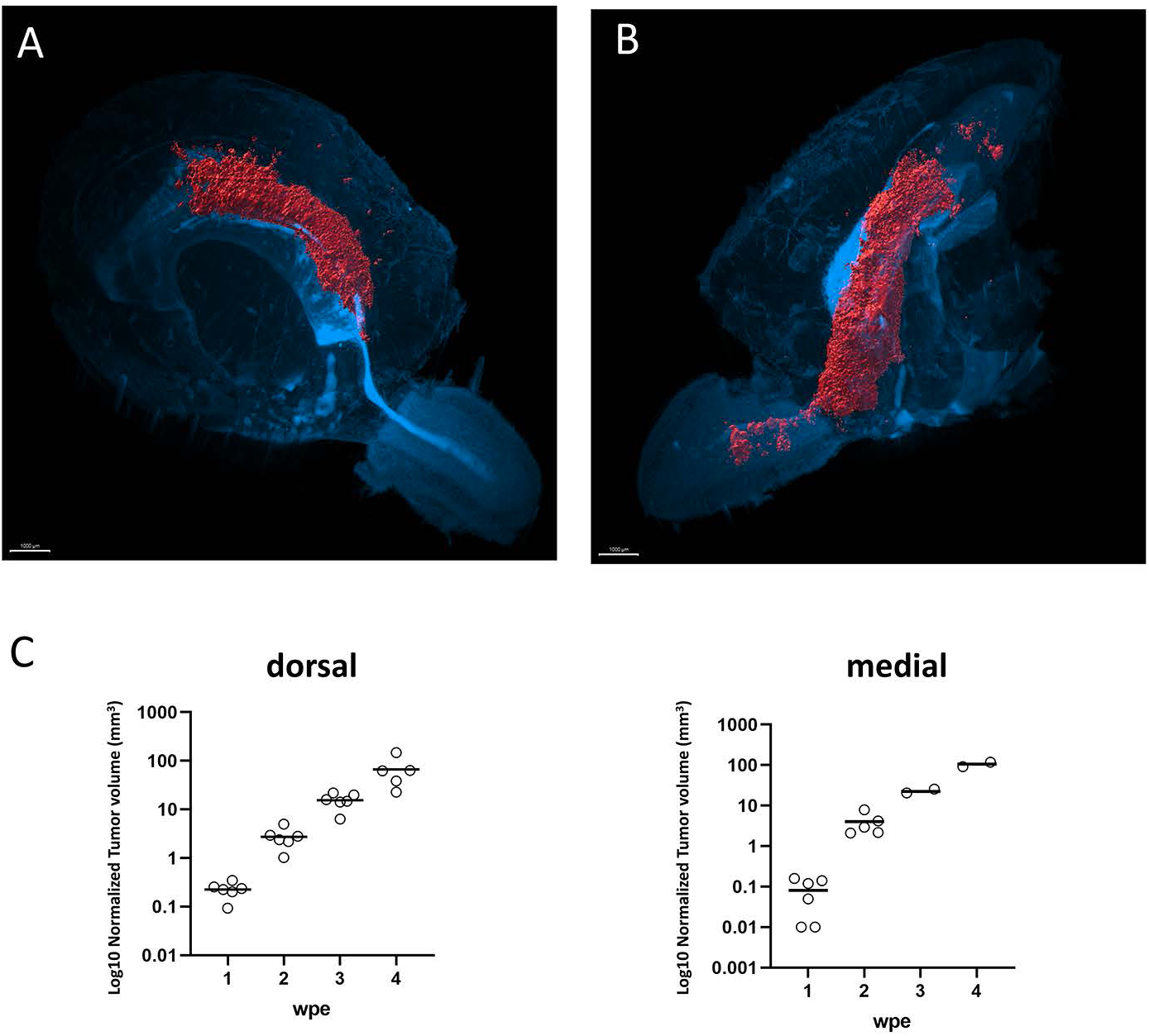
3D imaging of tumor at 2 weeks after electroporation targeting the dorsal wall (DWE; A) or the medial wall (MWE; B) of the lateral ventricle. (C) Graphs representing the growth of the tumor volume at 1, 2, 3 and 4 wpe in animals after DWE or MWE. Each circle represents one animal. The mean volume per time point per condition is drawn as a bar. Scale bars= 1000 µm.

**Supplementary S2.**
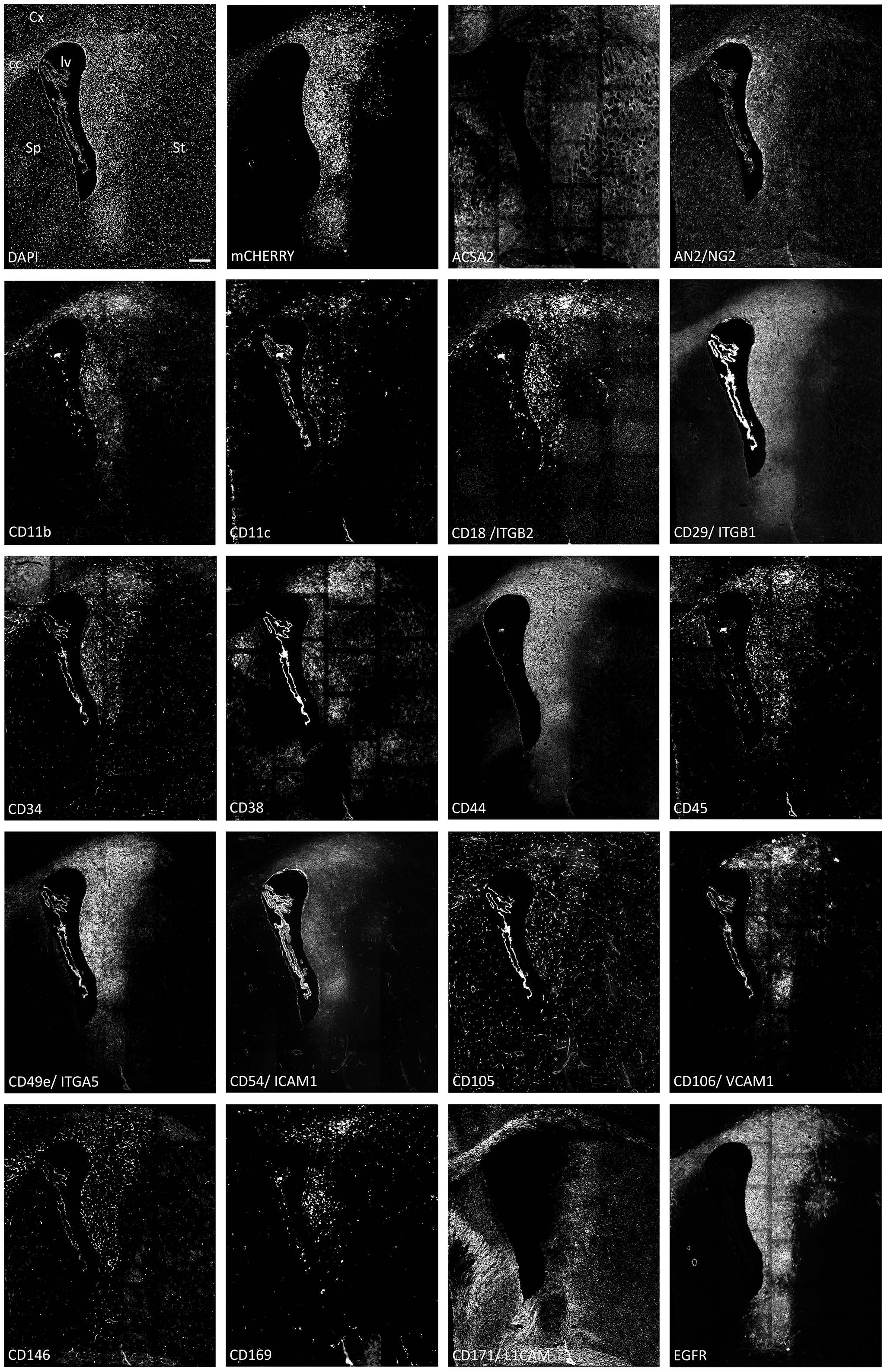
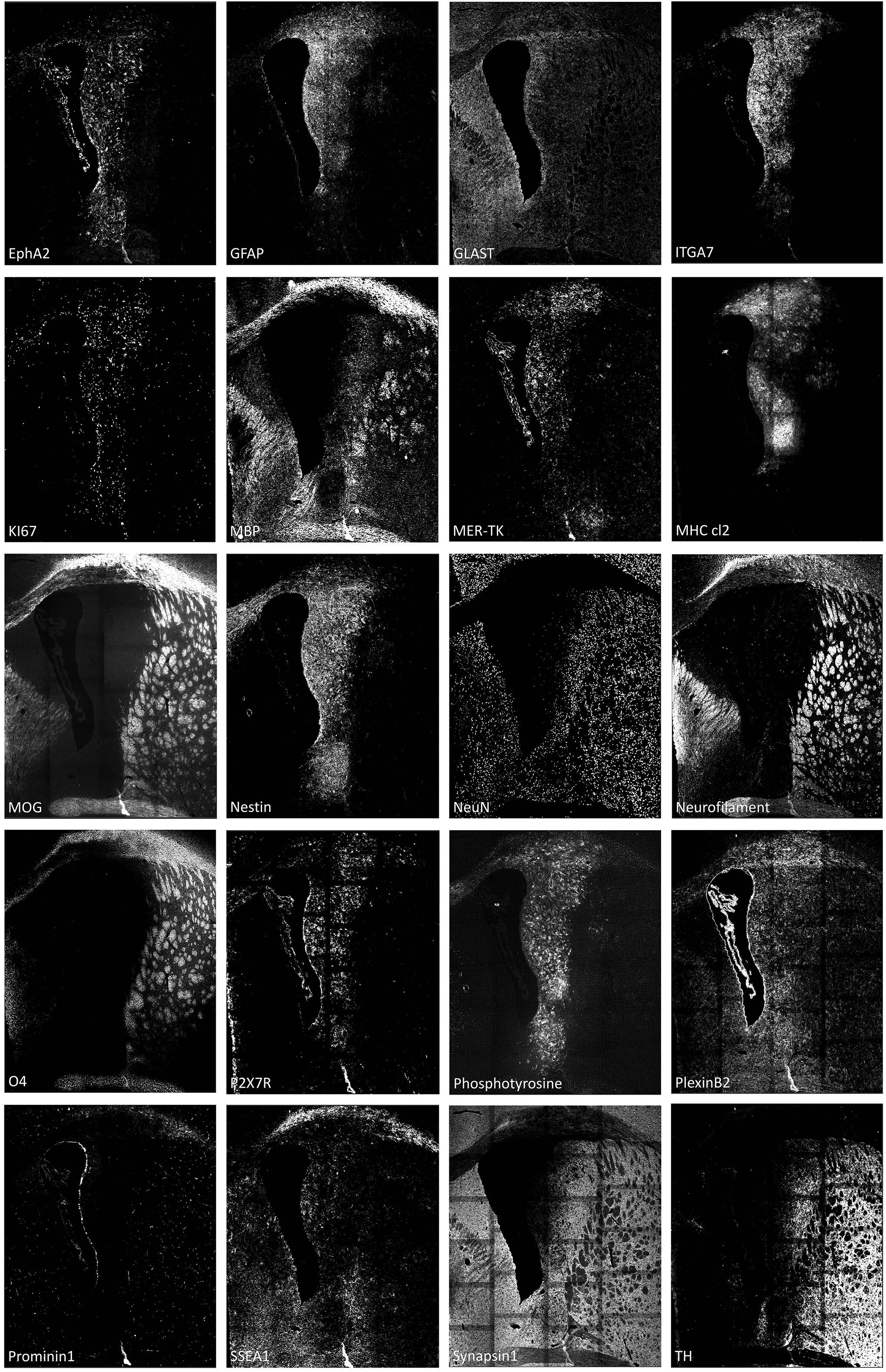
Single staining of DAPI, mCherry and each of the 38 selected antibodies used in MICS analyses on the selected coronal section at 2wpe after LWE. Scale bars= 250 µm. cc: corpus callosum; Cx: cortex; Sp: septum; St: striatum.

**Supplementary S3.**
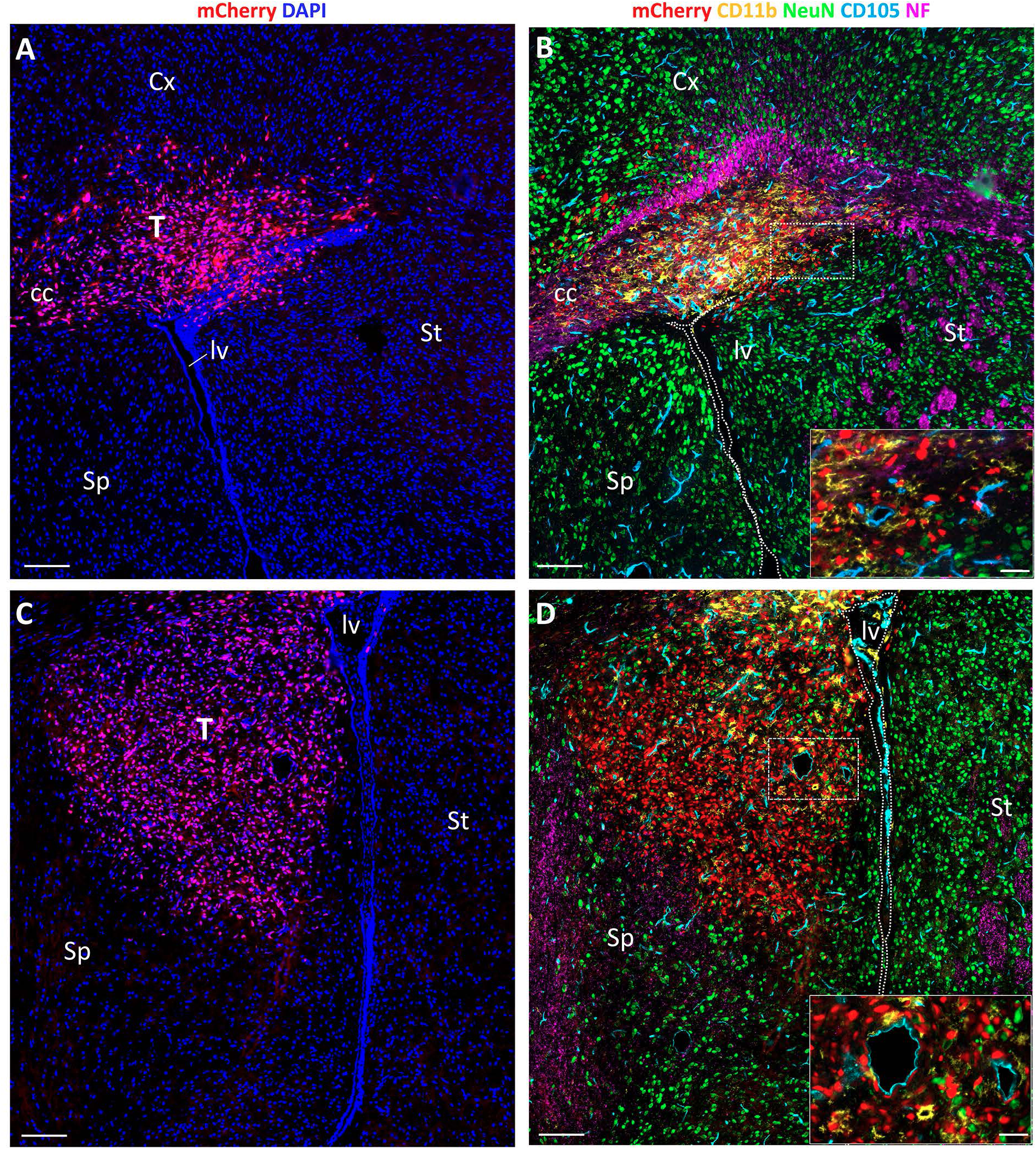
(A and C) Coronal sections used for MICS analyses showing mCherry positive cancer cells after DWE (A) and MWE (C). (B and D) Ultrahigh content imaging of these DWE (B) and MWE (D) sections with markers for tumor cells (mCherry), TAMs (CD11b), vascular endothelial cells (CD105), neurons (NeuN) and axonal tracts (NF; neurofilament). The boxed areas delineate high magnifications presented in the lower right corner, allowing the identification of the different cell types at cellular resolution. cc: corpus callosum; Cx: Cortex, lv: lateral ventricle, Sp: Septum; St: Striatum, T: Tumor. Scale bars= 500 µm (50 µm in insets)

**Supplementary S4.**
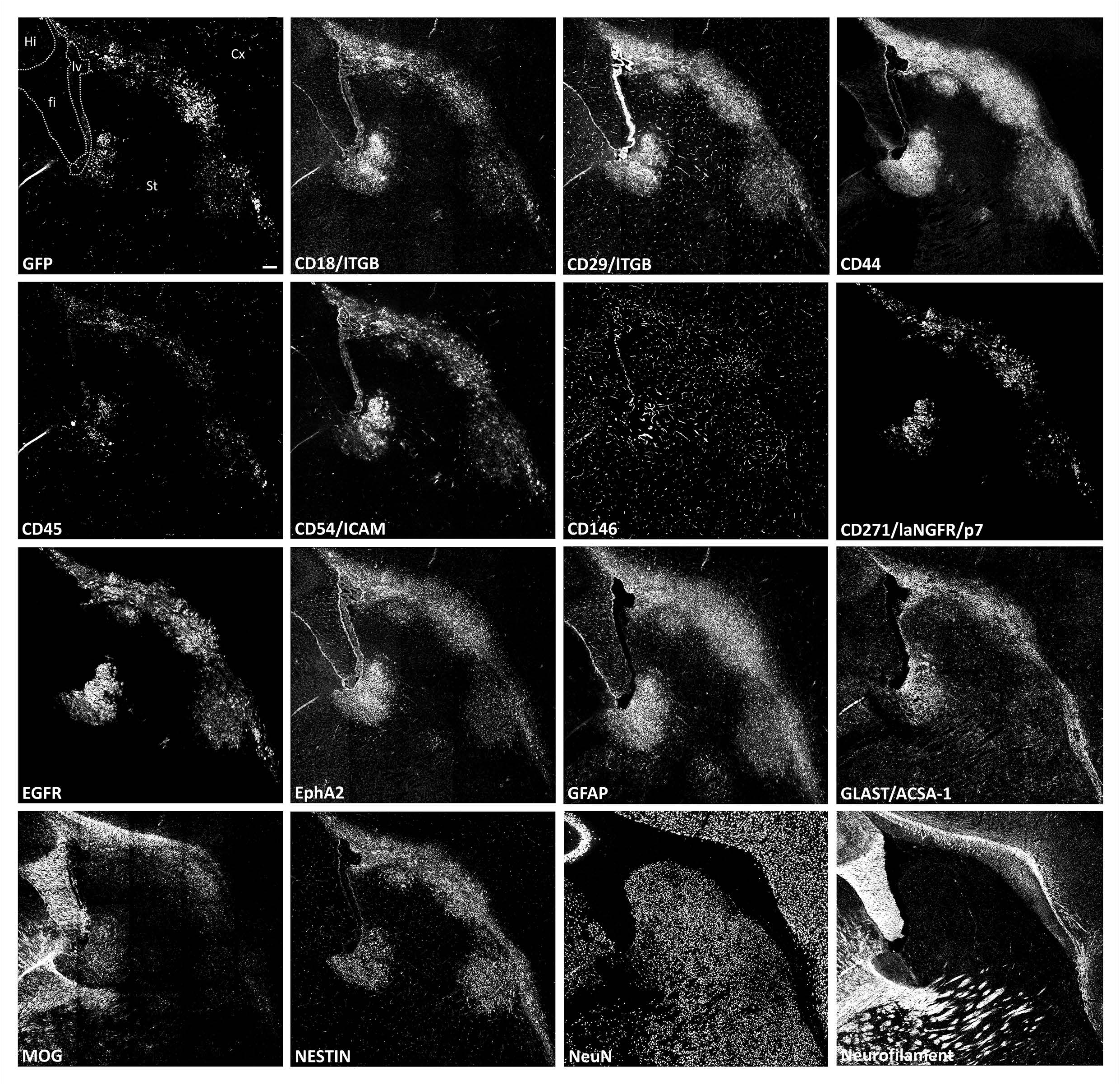
Individual staining patterns of antibodies functional in the 3D-2D approach as listed in Table 2. Scale bar= 100 µm. cc: corpus callosum, Cx: Cortex, fi: fimbria, Hi: Hippocampus, lv: lateral ventricle, St: Striatum.

